# Motor unit mechanisms of speed control in mouse locomotion

**DOI:** 10.1101/2024.12.29.628022

**Authors:** Kyle Thomas, Rhuna Gibbs, Hugo Marques, Megan R. Carey, Samuel J. Sober

## Abstract

During locomotion, the coordinated activity of dozens of muscles shapes the kinematic features of each stride, including systematic changes in limb movement across walking speed. Motor units, each of which consists of a single motor neuron and the muscle fibers it innervates, contribute to the total activation of each muscle through their recruitment and firing rate when active. However, it remains unknown how the nervous system controls locomotor speed by changing the firing of individual motor units. To address this, we combined quantitative behavioral analysis of mouse locomotion with single motor unit recordings from the lateral and long heads of the triceps brachii, which drive monoarticular extension of the elbow and biarticular movements of the elbow and shoulder, respectively. In contrast to prior studies employing bulk EMG to examine muscle activity, our recordings revealed the diversity of spike patterning across motor units as well as systematic differences in motor unit activity across muscles and locomotor speeds. First, motor unit activity differed significantly across the lateral and long heads, suggesting differential control of these two closely apposed elbow extensor muscles. Second, we found that individual units were recruited probabilistically (during only a subset of strides), showing that the highly repeatable bulk EMG signals observed across strides in fact reflect varying subsets of individual motor units. Finally, although recruitment probability and firing rate both increased at faster walking speeds, increases in recruitment were proportionally larger than rate changes, and recruitment of individual units accompanied changes in limb kinematics. Together, these results reveal how the firing of individual motor units varies systematically across muscles and walking speeds to produce flexible locomotor behavior.

## Introduction

Skilled behavior depends on the nervous system’s precise control of muscle activity. Motor units, which consist of a single motor neuron and all of the muscle fibers it innervates, generate the forces behind movement through their firing patterns. In locomotion, proper neural coordination of motor units within and across muscles allows for the stereotyped yet rapidly adjustable movement used for each step (Akay et al., 2014; Mayer & Akay, 2018; N. P. Schumann et al., 2006). In principle, the total force output of a muscle is modulated by the number of recruited motor units and the firing rate of active units (Enoka & Duchateau, 2017; Heckman & Enoka, 2012), with each newly-recruited unit increasing total muscle force by activating more muscle fibers. The firing rate and inter-spike-interval (ISI) pattern of recruited units then shape force production in concert with the biomechanics of the musculoskeletal system (Sober et al., 2018; Sponberg et al., 2011). Although studies in primates, cats, and zebrafish have shown that both the number of active motor units and motor unit firing rates increase at faster locomotor speeds (Grimby, 1984; Hoffer et al., 1981, 1987; Menelaou & McLean, 2012), the extent to which speed-dependent changes in rate and recruitment vary across muscles and species is unknown.

Mice demonstrate both physiological and biomechanical differences from other vertebrates, potentially leading to unique coordination among their motor units. Compared to cats, for example, mice have highly excitable motor units (Manuel et al., 2019; Manuel & Heckman, 2011) with muscle fibers heavily biased towards fast-twitch fibers (Burkholder et al., 1994; Mathewson et al., 2012), leading to rapid force production. Mice also locomote with greater stride frequency than larger species in order to achieve comparable speeds, requiring faster muscle activation and deactivation (Heglund & Taylor, 1988; Machado et al., 2015). The capability and need for faster force generation during dynamic behavior could implicate motor unit recruitment as a primary mechanism for modulating force output in mice (Manuel & Heckman, 2011; Dideriksen et al., 2020).

To quantify the organization of motor unit firing patterns during locomotion, we recorded mouse motor unit activity from the long head and lateral head of the triceps brachii during treadmill walking at various speeds. Both muscles extend the elbow while the long head also extends, rotates, and abducts the shoulder (Tata Ramalingasetty et al., 2021). Although bulk EMG recordings have shown that the triceps brachii is active in every step during quadrupedal locomotion (English, 1978; N. Schumann, 2002; Kirk et al., 2024), it is unknown how individual motor units are coordinated to generate this rhythmic pattern and whether motor pools from closely apposed muscles exhibit the same coordination. Using Myomatrix electrodes (Chung et al., 2023; Gilmer et al., 2024; Kirk et al., 2024) to record populations of individual motor units during locomotion, we found that units were recruited probabilistically across strides. When active, units fired in distinct locomotor phases with systematic differences in spike patterns across the long and lateral heads. At faster walking speeds, motor units increased both their recruitment probabilities and (to a lesser extent) their firing rates. Moreover, motor unit recruitment was correlated with variations in limb kinematics both within and across locomotor speeds, with recruitment of long head and lateral head units associated with different changes in limb movement. Overall, our results reveal the systematic changes in motor unit firing that regulate locomotor speed.

## Results

We collected kinematic and high-resolution electromyographic (EMG) data from six mice walking on a transparent treadmill that provided simultaneous lateral and ventral views of the animal (Darmohray et al., 2019). Using DeepLabCut (Mathis et al., 2018) to track body movements during locomotion (Figure 1A), we identified the stance phase of the right forelimb, defined as the period between footstrike and liftoff of the right forepaw (Figure 1B). Additionally, computing the internal angle of the elbow joint revealed that the elbow was minimally extended approximately 50 milliseconds before the footstrike (blue squares, Figure 1C). Electrode arrays (32-electrode Myomatrix array model RF-4x8-BHS-5) were implanted in forelimb muscles (note that Figure 1D shows the EMG signal from only one of the 16 bipolar recording channels), and the resulting data were used to identify the spike times of individual motor units in the triceps brachii long and lateral heads (Table 1, Figure 1E) as described previously (Chung et al., 2023). To best capture the spike pattern given that some units begin firing prior to footstrike, we defined a stride cycle as the period between minimum elbow angles rather than consecutive footstrikes. Kinematic analysis of locomotor data at different walking speeds revealed systematic variation in the temporal (Figure 1F) and spatial (Figure 1G, H) components of limb movement, consistent with prior reports (Akay et al., 2006; Bellardita & Kiehn, 2015; Machado et al., 2015; Mendes et al., 2015).

**Figure 1.**
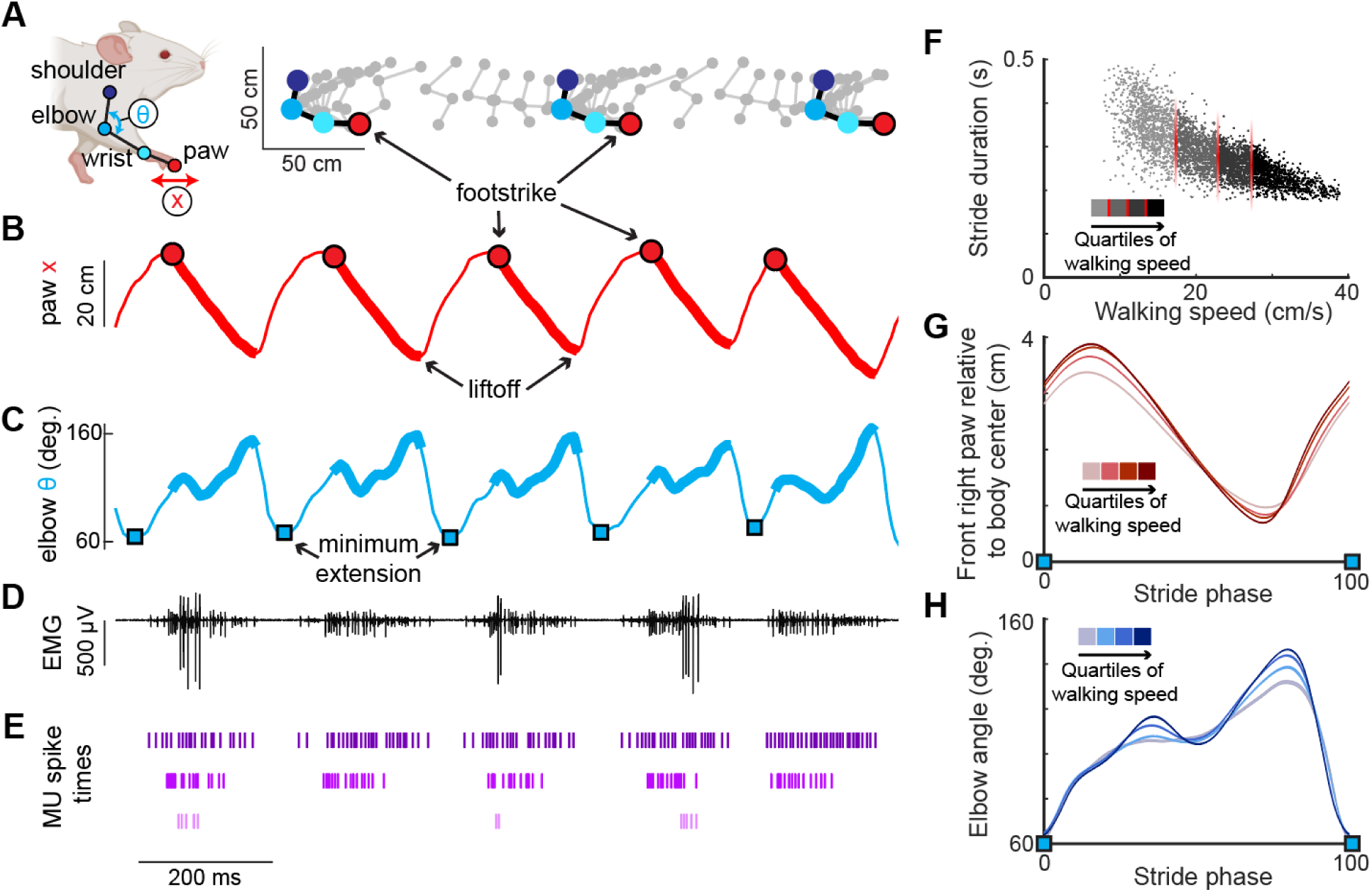
High-resolution muscle recording during mouse locomotion. **(A)** (Left) Anatomical landmarks (shoulder, elbow, wrist, paw) and kinematic features (elbow angle θ, paw position x) tracked via high-resolution video during treadmill walking (Darmohray et al., 2019; Mathis et al., 2018). (Right) Position of anatomical landmarks during two stride cycles with limb position captured every 15 ms. **(B)** Position of the right forepaw (x) relative to body center. Thick lines represent the stance phase when the paw is in contact with the floor of the treadmill. **(C)** Interior elbow angle (θ) during locomotion. Troughs of this measure, denoting minimum extension (blue squares), were used to define the spike window for each stride. **(D)** Representative single channel of electromyographic (EMG) activity recorded from the long head of the right triceps. **(E)** Three motor units recorded from the long head identified from the above EMG trace (see Methods). Note that the bottom-most unit is active in only a subset of stride cycles. **(F)** Relationship between stride duration and walking speed for all strides in an example mouse. Each dot represents a stride, with shading indicating the speed quartile within which the stride falls (see Methods). **(G,H)** Right forepaw position x (G) and elbow angle (H) within the walking speed quartiles. Both (G) and (H) are normalized to total stride duration beginning and ending with elbow minimum extension (blue squares) and show mean (± SE).

**Table 1.** Motor units identified per muscle in each experimental mouse. Each thread consisted of 8 electrode contacts used to record bipolar EMG, and numbers in parentheses indicate the number of motor units isolated using the spike sorting algorithm described in Methods. Data from biceps muscle implantation in two mice were not spike sorted and are not included in this report.

| Mouse | Implanted muscle (# motor units identified) |
| --- | --- |
| A | 2 threads right **lateral head, triceps brachii** (3)<br>2 threads right **long head, triceps brachii** (1) |
| B | 2 threads right **lateral head, triceps brachii** (0)<br>2 threads right **long head, triceps brachii** (3) |
| C | 2 threads right **lateral head, triceps brachii** (0)<br>2 threads right **long head, triceps brachii** (6) |
| D | 2 threads right **lateral head, triceps brachii** (4)<br>2 threads right biceps brachii (n/a) |
| E | 2 threads left biceps brachii (n/a)<br>2 threads right **long head, triceps brachii** (7) |
| F | 4 threads right **lateral head triceps brachii** (9) |

### Motor units are probabilistically recruited across strides

Despite the triceps muscles as a whole being reliably activated on every step (English, 1978; N. Schumann, 2002; Kirk et al., 2024), the majority of individual motor units in both the long head and lateral head were active only in a subset of strides during locomotion. Motor units in both muscles exhibited this pattern of probabilistic recruitment (defined as a unit’s firing on only a fraction of strides), but with differing distributions of firing properties across the long and lateral heads (Figure 2). For each motor unit, we measured the probability of a unit being recruited as the percentage of strides with at least one spike. Units demonstrated a variety of firing patterns, with some units producing 0 spikes more frequently than any non-zero spike count (Figure 2A, B), and units that were less likely to be recruited also had lower average spike counts (Figure 2–figure supplement 1). A subset of units, primarily in the long head, were recruited in under 50% of the total strides and with lower spike counts (Figure 2C). This distribution of recruitment probabilities might reflect a functionally different subpopulation of units. However, the distribution of recruitment probabilities were not found to be significantly multimodal (p>0.05 in both cases, Hartigan’s dip test; Hartigan, 1985). However, Hartigan’s test and similar statistical methods have poor statistical power for the small sample sizes (n=17 and 16 for long and lateral heads, respectively) considered here, so the failure to achieve statistical significance might reflect either the absence of a true difference or a lack of statistical resolution.

**Figure 2.**
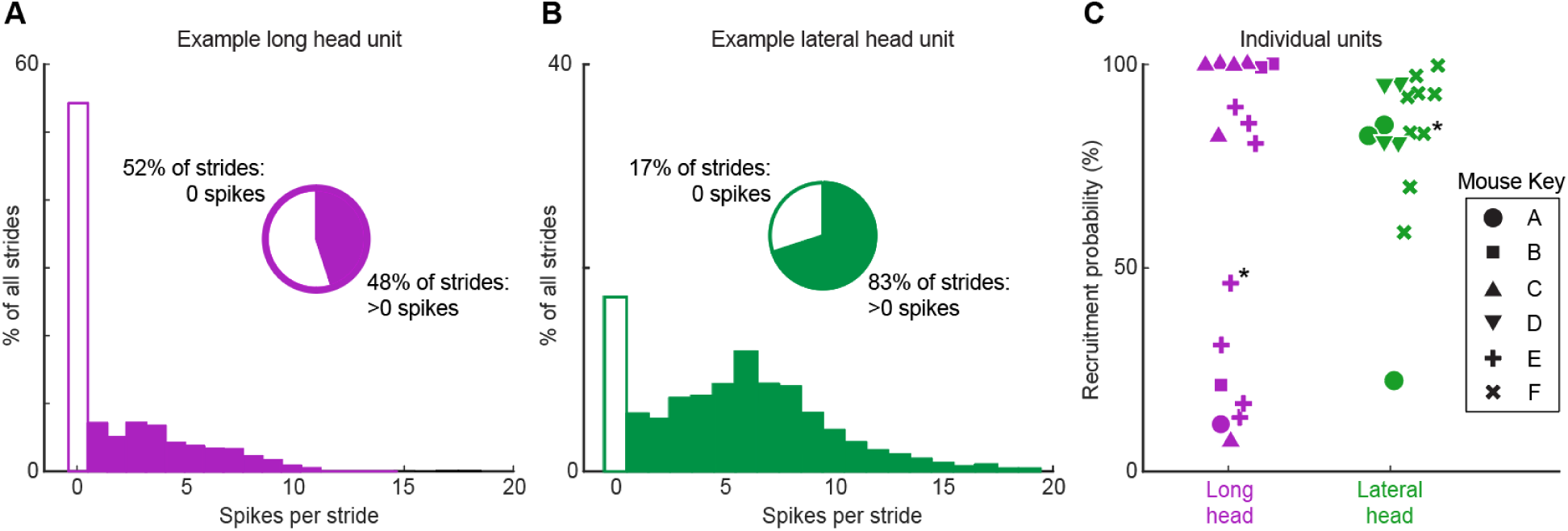
Motor unit spike count distributions. **(A)** Example motor unit from the long head of the triceps muscle fired zero spikes on 52% of strides, but on the other 48% of strides fired 1-14 spikes. **(B)** Example motor unit from the triceps lateral head fired zero spikes in 17% of strides but 1-19 spikes on the other 83% of strides. **(C)** Percentage of strides with at least one spike (probability of recruitment) for all recorded motor units in the long (purple) and lateral (green) heads of the triceps. Symbols denote different animals and each point reflects an individual motor unit. Asterisks highlight the units shown in (A,B).

### Motor unit firing patterns in the long and lateral heads of the triceps

Motor units within each muscle fired at distinct phases of the stride cycle. Units in the long head typically became active near the time of footstrike, with approximately half of the units reliably recruited prior to footstrike (Figure 3A,B). In contrast, units in the lateral head began spiking after the long head was already active and remained active until just prior to liftoff (Figure 3B,C). Furthermore, units in the long head reached their stride-dependent peak rates before the lateral head (p<0.01, two-sample k-s test). These findings demonstrate that despite the synergistic (extensor) function of the long and lateral heads of the triceps at the elbow, the motor pool for the long head becomes active roughly 100 ms before the motor pool supplying the lateral head during locomotion (Figure 3C). This timing difference suggests distinct patterns of synaptic input onto motor neurons innervating the lateral and long heads. In contrast to the timing differences described above, motor units in the lateral and long heads displayed similar burst durations (Figure 3B,E) and peak firing rates (Figure 3D).

**Figure 3.**
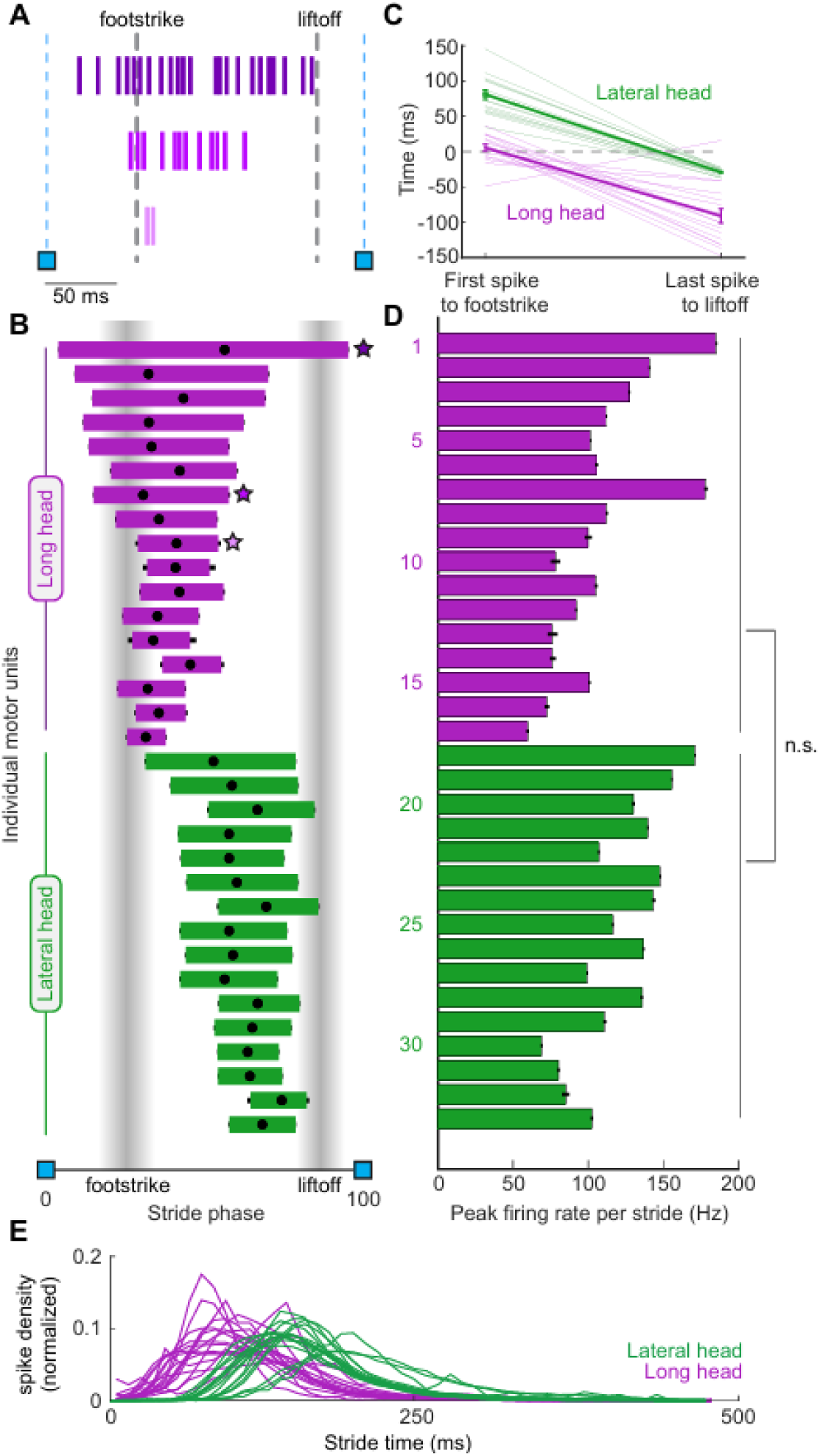
Motor unit firing patterns within and across muscles. **(A)** Example stride with three units from the long head. **(B)** Mean phase (± SE) of motor unit burst activity within each stride duration across all strides. Black dots on each bar show the mean phase of the unit’s peak firing rate. Starred points refer to the examples in A. **(C)** Left: Mean time (± SE) between the first spike of a unit’s spike train and the right forepaw footstrike. Positive values denote the spike happening after the footstrike. Right: Mean time (± SE) between the last spike of a unit and the liftoff. Light traces denote values for individual motor units, the heavy trace shows the mean (± SE) across all units within a muscle. **(D)** Mean peak firing rate (± SE) of each unit. Note that these measurements only include strides in which the given unit was recruited. **(E)** Spike probability density. Each trace shows the probability density function of all spikes recorded from each motor unit. Stride times in (E) are aligned such that time=0 represents the time of minimum elbow angle (Fig. 1C).

The evolution of spike patterns within each stride differed between motor unit populations in the long and lateral heads. In both muscles, motor units with longer burst durations reached higher peak firing rates (Figure 4A). However, the slope of this relationship was significantly higher for lateral head units (p<0.05, permutation test). We also observed muscle-dependent differences in motor unit patterning when examining the inter-spike intervals (ISIs) between the first three spikes in each stride cycle. Motor units in both muscles began firing with ISIs typically below 12 ms (Figure 4B). Furthermore, the second ISI was generally shorter, indicating that firing rate increased throughout the first three spikes fired in the stride cycle. However, the population of motor units in the long head had a larger magnitude in the ratio ISI_1_/ISI_2_ (Figure 4B, p<0.01, two-sample k-s test). Together with the differences in burst timing shown in Figure 3B, these results again suggest that the motor pools for the lateral and long heads of the triceps receive distinct patterns of synaptic input, although differences in the intrinsic physiological properties of motor neurons innervating the two muscles might also play an important role.

**Figure 4.**
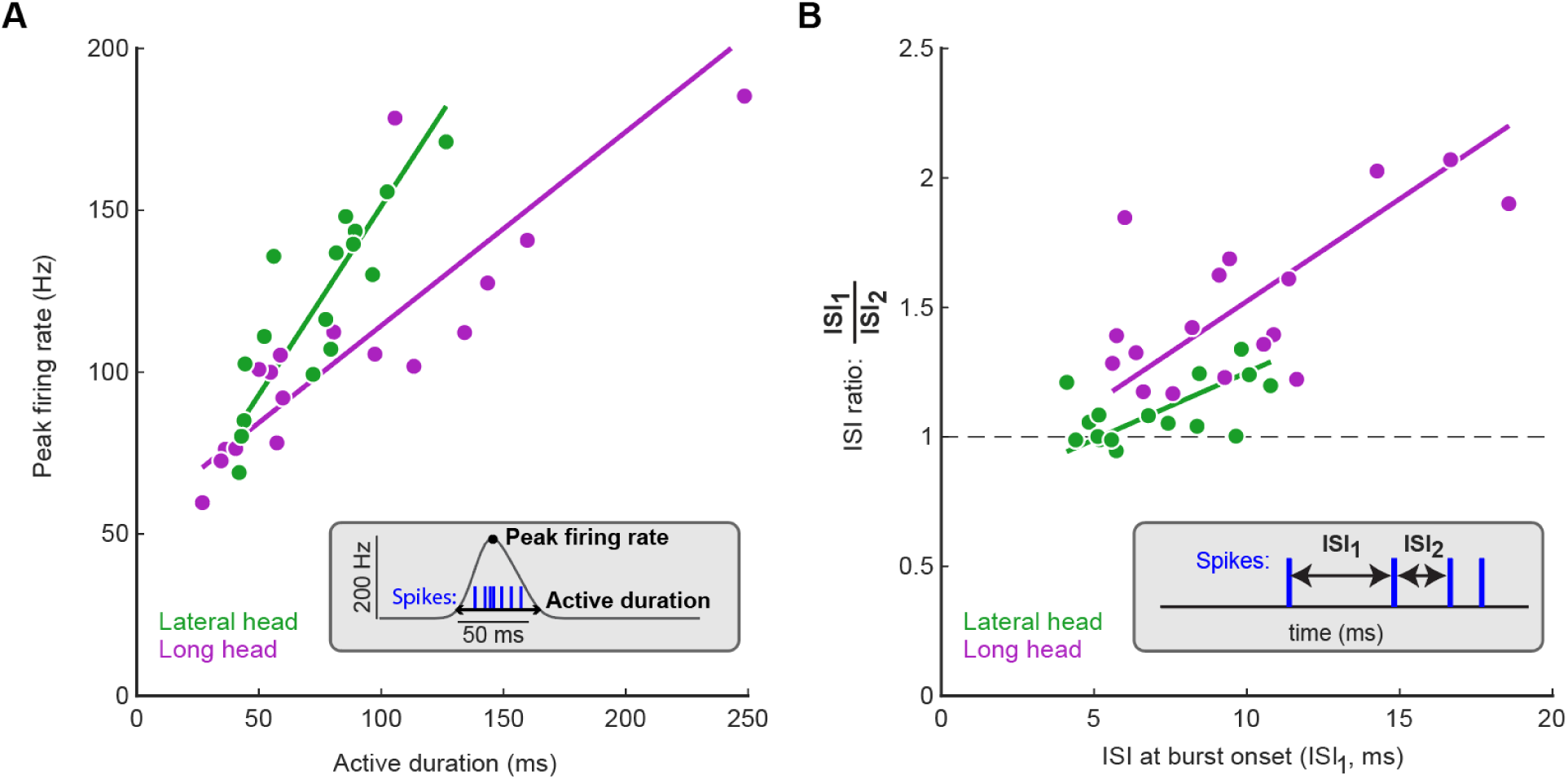
Motor unit spike patterns evolve differently in the long and lateral heads. **(A)** Relationships between active duration and peak firing rate across motor pools. Type 2 regression slopes were significantly different between the lateral and long heads (p<0.05, permutation test). **(B)** Motor unit inter-spike intervals (ISIs) across the first three spikes in motor unit bursts. Each data point shows the mean of the first ISI and the ratio between the first and second ISIs for a single unit. Note that by definition only strides with at least three spikes could be used for the analysis shown in panel (B). Type 2 regression slopes were not significantly different between data from the lateral and long heads (p>0.05, permutation test). Data shown here are grouped across all mice; **Figure 4-figure supplement 1** shows how these data are distributed across animals.

### Motor unit mechanisms of speed control

Adjusting walking speed requires changes in the firing patterns of individual motor units, which could include speed-dependent changes in units’ probability of recruitment and/or changes in firing rate. To investigate the changes in motor unit firing underlying locomotor speed control, we quantified how both recruitment probability and firing rate change across the four quartiles of locomotor speed shown in Figure 1F-H. Motor units from the long and lateral heads of the triceps (Figure 5A,B, purple and green traces, respectively) displayed significant increases in recruitment probability as locomotor speeds increased. Figure 5C shows each motor unit’s difference in recruitment probability between the slowest and fastest locomotor speed quartiles. This increase was statistically significant in 31/33 motor units in our study (p<0.05, permutation test) when considered individually, and was also significant when the probabilities of all motor units were analyzed as a group (p<0.01, Wilcoxon signed-rank test). Robust increases in recruitment probability across the four speed quartiles were therefore the norm in our dataset.

**Figure 5.**
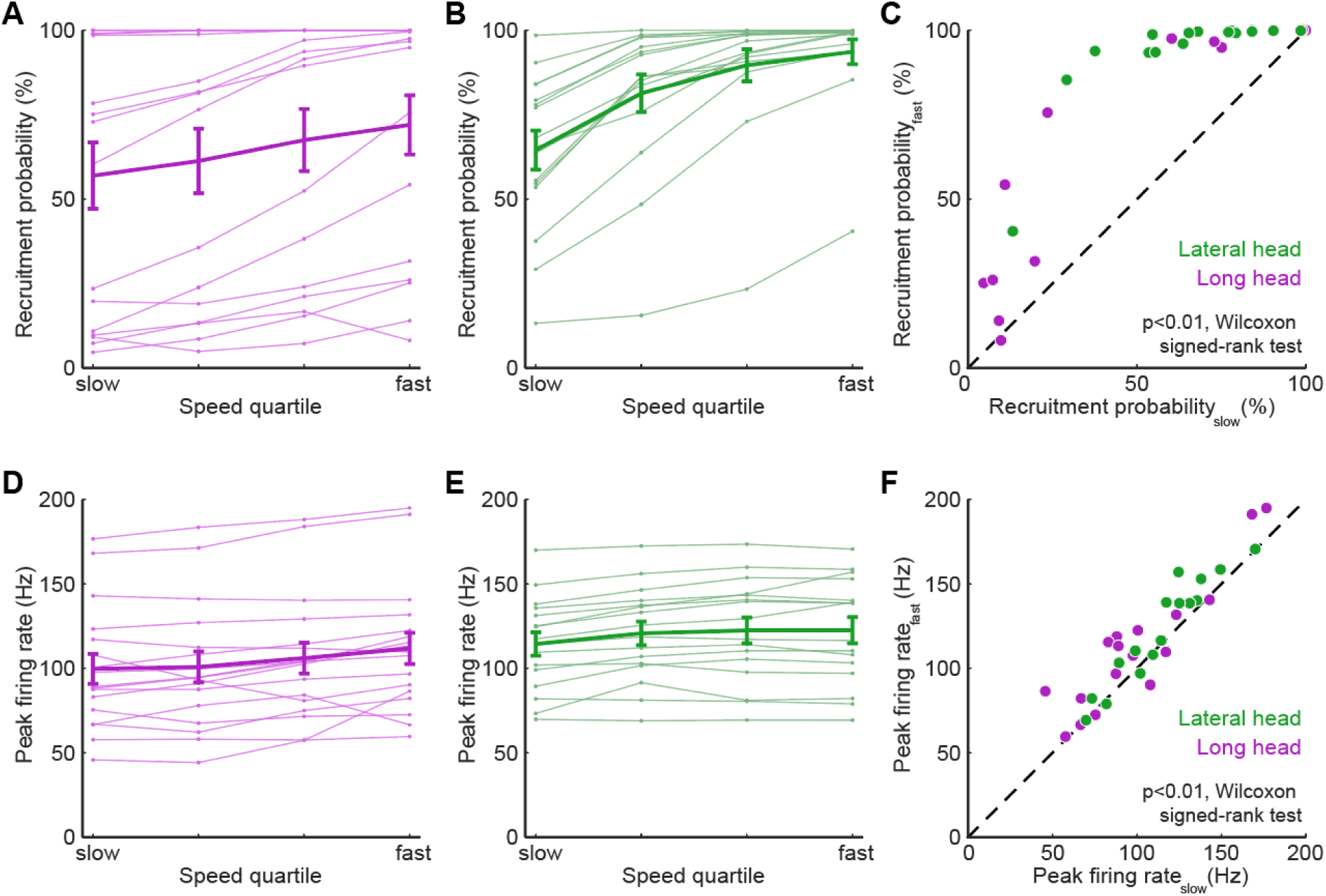
Motor units alter firing rate and recruitment across walking speeds. **(A)** Light traces show median of recruitment probability for individual long head motor units while the heavy trace shows mean (± SE) across all long head motor units. (**B)** Recruitment probability for lateral head motor units, same plotting conventions as in (A). **(C)** Difference in recruitment probabilities between slowest and fastest speed quartiles for all motor units. **(D)** Light traces show median of peak firing rate for individual long head motor units while the heavy trace shows mean (± SE) across all long head motor units. (**E)** Peak firing rates for lateral head motor units, same plotting conventions as in (D). **(F)** Difference in peak firing rates between slowest and fastest speed quartiles for all motor units. Across all motor units, both recruitment probabilities (C) and firing rates (F) were significantly higher at the fastest quartile than at the slowest quartile (p<0.01, Wilcoxon signed-rank tests). Speed-dependent changes in recruitment vs speed-dependent changes in firing rate for individual motor units and experimental animals are shown in **Figure 5–figure supplement 1** and **Figure 5–figure supplement 2**, respectively.

Quantitative analysis of motor unit activity also revealed significant speed-dependent changes in firing rate, although these were proportionally smaller than the increases in recruitment probability. Motor units in both the long and lateral heads of the triceps (Figure 5D,E, purple and green traces, respectively) often had either marginal increases or no difference in peak firing rate at faster speeds. Across all motor units in our dataset in the slowest and fastest speed quartiles (Figure 5F), we observed significant increases in peak firing rate in 22/33 individual motor units in our study (p<0.05, permutation test), and also a significant speed-dependent increase in peak rate when considering all motor units together (p<0.01, Wilcoxon signed-rank test). Speed-dependent increases in peak firing rate were therefore also present in our dataset, although in a smaller fraction of motor units (22/33) than changes in recruitment probability (31/33). Furthermore, the mean (± SE) magnitude of speed-dependent increases was smaller for spike rates (mean rate_fast_/rate_slow_ of 111% ± 20% across all motor units) than for recruitment probabilities (mean p(recruitment)_fast_/p(recruitment)_slow_ of 179% ± 3% across all motor units). While fractional changes in rate and recruitment probability are not readily comparable given their different upper limits, these findings could suggest that while both recruitment and peak rate change across speed quartiles, increased recruitment probability may play a larger role in driving changes in locomotor speed.

### Kinematic contributions of motor unit recruitment

We next examined whether the probabilistic recruitment of individual motor units in the triceps – an elbow extensor muscle – was correlated with stride-by-stride variations in elbow angle kinematics. To do so, we compared elbow extension (Δθ; Figure 6A) on strides in which each individual motor unit did or did not fire at least one spike. When kinematic data are combined across all speed quartiles (Figure 6B), we found that recruitment of lateral head motor units (green symbols) is associated with greater elbow extension, whereas recruitment of long head units (purple symbols) is correlated with smaller extensions (p<0.001, Wilcoxon signed-rank tests). These correlations might reflect both an influence of motor unit recruitment on limb kinematics as well as different biomechanical roles for the long and lateral heads.

**Figure 6.**
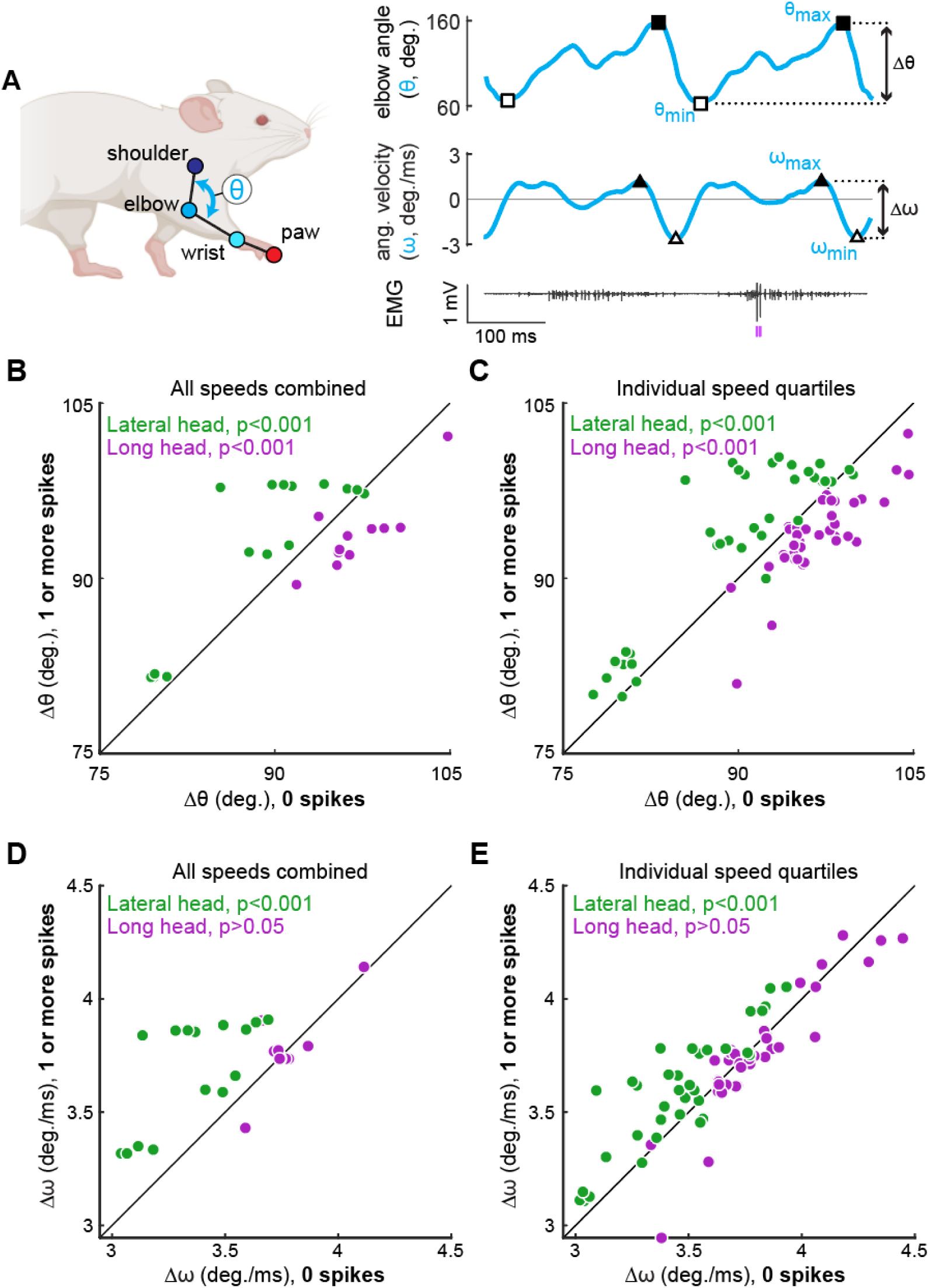
Motor unit recruitment correlates with muscle-specific kinematic differences. **(A)** We calculated the ranges of elbow angle (Δθ) and elbow velocity (Δω) observed on strides in which each motor unit did or did not fire at least one spike (purple tick marks below EMG trace). **(B,D)** Each point represents the mean Δθ (B) or Δω (D) observed on strides in which a single motor unit fires zero spikes (horizontal axis) vs. when the motor unit fires at least one spike (vertical axis). Note that in these panels, data for each motor unit were combined across all locomotor speeds. **(C,E)** Same analyses as before, except each motor unit contributes up to four data points, one for each of the four locomotor speed quartiles in which sufficient data were available (at least 30 strides existed in both the spiking and non-spiking conditions within a given quartile). Legends in each panel show statistical significance for a difference in kinematics tested on the motor units within each muscle (Wilcoxon signed-rank tests). Note that most of the muscle-specific differences shown in (C,E) were also present when each of the four quartiles (**Figure 6-figure supplement 1**) or experimental animals (**Figure 6-figure supplement 2**) were examined individually.

Since both limb kinematics (Figure 1G,H) and recruitment probability (Figure 5) are significantly correlated with locomotor speed, the observed correlations between unit recruitment and elbow angle across all speeds (Figure 6B) does not necessarily reveal the direct influence of unit firing on limb kinematics. We therefore controlled for speed by repeating the analysis shown in Figure 6B for strides within each speed quartile. Strikingly, the correlations between motor unit recruitment and elbow angle persisted in this alternative analysis (Figure 6C; p<0.001, Wilcoxon signed-rank tests), suggesting that the recruitment of individual motor units in the lateral and long heads might have significant effects on elbow angle in strides of similar speed (see Discussion). We repeated these analyses using the elbow angular velocity rather than just the angle to further identify how firing patterns related to behavior. Motor units in the lateral head had a similar effect with larger velocities correlating with motor unit recruitment across all speeds (Figure 6D,E; p<0.001).

### Probabilistic recruitment is correlated across motor units

Our results show that the recruitment of individual motor units is probabilistic even within a single speed quartile (Figure 5A-C) and predicts body movements (Figure 6), raising the question of whether the recruitment of individual motor units are correlated or independent. Correlated recruitment might reflect shared input onto the population of motor units innervating the muscle (De Luca, 1985; De Luca & Erim, 1994; Farina et al., 2014). For example, two motor units, each with low recruitment probabilities, may still fire during the same set of strides. To assess the independence of motor unit recruitment across the recorded population, we compared each unit’s empirical recruitment probability across all strides to its conditional recruitment probability during strides in which another motor unit from the same muscle was recruited (Figure 7). Doing this for all motor unit pairs revealed that motor units in both muscles were biased towards greater recruitment when additional units were active (p<0.001, Wilcoxon signed-rank tests for both the lateral and long heads of triceps). This finding suggests that probabilistic recruitment reflects common synaptic inputs that covary together across locomotor strides.

**Figure 7.**
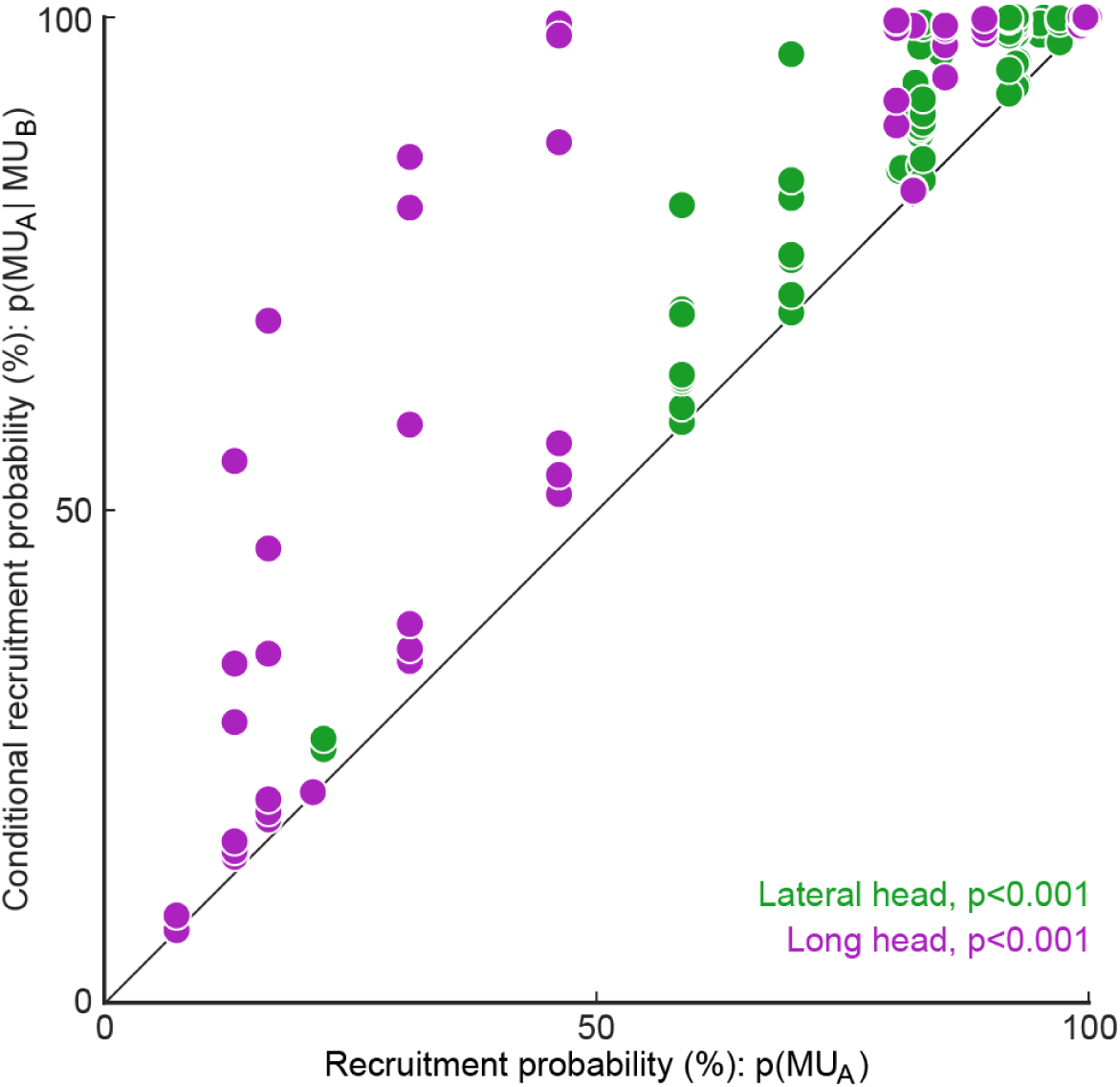
Motor unit recruitment probability is greater when other motor units within the muscle are recruited. Each point reflects a motor unit’s empirically measured recruitment probability, *p(MU_A_),* across all strides compared to strides when a simultaneously-recorded motor unit from the same muscle was recruited, *p(MU_A_|MU_B_)*. Motor units in each muscle had significantly higher recruitment when another unit was recruited in the same stride (p<0.001, Wilcoxon signed-rank tests).

## Discussion

Our results quantify the heterogeneity of individual motor units’ firing patterns across muscles and walking speeds. Motor units were probabilistically recruited on a stride-by-stride basis with peak firing rates between 60-185 Hz when active. Motor units in the long head were recruited before the lateral head and spiking patterns evolved within each stride differently across the two muscles. Motor units in both muscles demonstrated increases in their recruitment probability and firing rate at faster walking speeds. Furthermore, motor unit recruitment was also correlated with differences in limb kinematics for strides of similar speed. As discussed below, firing patterns from motor units in the long and lateral heads likely reflect the functional and anatomical role of these two muscles, highlighting the need for high-resolution quantification of motor unit firing patterns during behavior.

### Differences in motor unit activity patterns across two elbow extensors

Motor unit spike patterns differed systematically between the long and lateral heads of the triceps brachii. Motor units in the long head were consistently recruited earlier than units in the lateral head (Figure 3B,C). The large differences in burst timing and spike patterning across the muscle heads suggest that the motor pools for each muscle receive distinct inputs. However, differences in the intrinsic physiological properties of motor units and neuromodulatory inputs across motor pools might also make substantial contributions to the structure of motor unit spike patterns (Martínez-Silva et al., 2018; Miles & Sillar, 2011).

The observed order of muscle activation matches past reports of bulk muscle activity in these two muscles across other quadrupedal species (Carroll & Biewener, 2009; Drew et al., 2008; Livingston & Nichols, 2014; Scholle et al., 2001) and may reflect the biomechanical functions of each muscle. Whereas the lateral head is a monoarticular elbow extensor, the long head is biarticular, both extending the elbow along with extending and rotating the shoulder (Tata Ramalingasetty et al., 2021). Although we did not measure ground reaction forces in this study, prior reports indicate that the vertical ground reaction force on the mouse forepaw reaches two peaks during locomotion (Schmitt et al., 2010). The first peak, which happens soon after the footstrike, has a lower magnitude than the second peak, which occurs closer to liftoff (Clarke et al., 2001; Schmitt et al., 2010). Studies in both rats (Sarver et al., 2010) and cats (Corbee et al., 2014) have demonstrated that horizontal ground reaction forces in both the medio-lateral and cranio-caudal directions are also greatest soon after footstrike, with more force variability than the vertical reaction force. Since units in the long head are most active following footstrike, this suggests that activity in the long head might be related to stabilizing the limb within each step. Our finding that recruitment of long head motor units (purple symbols, Figure 6) accompanied smaller elbow extensions might therefore reflect a more complex biomechanical role for the long head, potentially relating to shoulder rotation (which was not measured in this study). This interpretation is consistent with past findings that biarticular muscles are power distributors, stabilizing the joint across multiple dimensions, while monoarticular muscles are power generators (Ryan & Gregor, 1992; Van Ingen Schenau et al., 1992, 1994). Along these lines, the observed timing of lateral head motor unit activity just prior to liftoff (Figure 3B) might therefore reflect the lateral head’s role of providing propulsion prior to swing, consistent with our finding that recruitment of motor units in the lateral head is correlated with both larger elbow extension and more rapid changes in angle (Figure 6).

Motor units in the lateral and long heads also differed with respect to their recruitment probabilities, with a substantial population of units in the long head (but not the lateral head) with probability of recruitment less than 50% (Figure 2C). This difference may reflect different functions of muscle fibers in different subcompartments of biarticular muscles. Prior work has established that different regions within a biarticular muscle can have different contributions across the two joints (Chanaud et al., 1991; English & Weeks, 1987; Watanabe et al., 2021). For example, different regions of the cat biceps femoris are out of phase with each other during walking, with the anterior compartment active during stance as a hip extensor and the posterior compartment active during swing as a knee flexor (Chanaud et al., 1991; English & Weeks, 1987). Additionally, the posterior compartment was only active at faster speeds. In our mouse data, functional compartments within the biarticular long head might similarly explain the differently recruited populations for motor units (Figure 2C). However, to our knowledge, no studies have investigated anatomical or functional subdivisions across subregions of the triceps long head in the mouse. Nevertheless, the group of less-frequently recruited units might contribute more to forelimb joint stability during a small number of strides affected by rare and unexpected external perturbations whereas the other long head motor units might be recruited in a greater fraction of strides to support the weight of the body. Further examination of the anatomical microstructure of the long head, including precise characterization of the attachment points to the bone (DeWolf et al., 2024; Gilmer et al., 2024), are necessary to answer these questions.

The varied composition of fiber types in the long and lateral heads may also explain the different firing patterns across muscles. Although both muscles are heavily biased towards the fastest myosin type (type 2B), the long head has a broader composition, including a small percentage of slower isoforms as well (type 1 and 2A) (Mathewson et al., 2012). Type 2B isoforms are related to fast-twitch, fatigable units (FF) while type 1 compose slow-twitch units (S) and type 2A are intermediate between fast and slow (Bączyk et al., 2022; Schiaffino & Reggiani, 2011). While we were unable to directly quantify the unit type, the majority of units observed, particularly within the lateral head, are likely FF units given these prior histological findings. Thus, while we did not explicitly measure muscle fatigue after our recordings (up to 30 minutes of walking at 12.5-27.5 cm/sec, see Methods), it is possible that fatigue might have contributed to the observed probabilistic recruitment of later-recruited units (Martínez-Silva et al., 2018). Similarly, motor units that were recruited in nearly every stride with 10 or more spikes per stride (Figure 2–figure supplement 1) could represent a population with slower isoforms given their resistance to fatigue. The most prominent example of this came from the single unit in the long head that fired for over 90% of the stride phase in every stride (Figure 3B, top).

### Firing rates in mouse locomotion compared to other species

The range of firing rates we observed in mice are higher than those typically observed in larger species, likely reflecting the distinctive physiology of mouse motor neurons. Motor units in the lateral and long heads of the triceps exhibited a large and overlapping range of peak firing rates ranging from 50-175 Hz (Figure 3D), in agreement with prior reports of motor unit firing rates from mouse forelimb (Kirk et al., 2024) and hindlimb (Hadzipasic et al., 2016) during locomotion. In rat hindlimb muscles during walking, motor units had mean instantaneous firing rates between 45-109 Hz (Gorassini et al., 2000). During locomotion in cats, motor units recorded from the toe and hindlimb had firing rates between 15-50 Hz (Hoffer et al., 1987; Zajac & Young, 1980). Human motor units in the short extensors of the toe fire at even lower rates (10-25 Hz) during walking (Grimby, 1984). Compared to these larger species, mice likely reach higher rates through the physiological properties of their motor neurons such as afterhyperpolarization (AHP), which influences how rapidly a neuron returns to baseline voltage after firing a spike. Although AHP durations vary across unit types, AHP durations in mice are approximately two and three times shorter than those in cats and humans respectively (Manuel et al., 2009, 2019; Meehan et al., 2010). Additionally, persistent inward currents (PICs), which amplify excitatory synaptic inputs (Binder et al., 2020; Heckman et al., 2005), might lead to disproportionately large gain in mouse motor neurons compared to other species (Huh et al., 2017; Manuel et al., 2019). Consequently, even mice performing quiet standing have motor unit firing rates reaching up to 68 Hz (Ritter et al., 2014). Our findings (Figure 4) highlight that even with the relatively high firing rates observed in mice, there are still significant changes in firing rate and recruitment probability across the spikes within bursts (Figure 4B) and across locomotor speeds (Figure 5F). Future studies should more carefully examine how these rapidly changing spiking patterns derive from both the statistics of synaptic inputs and intrinsic properties of motor neurons (Manuel & Heckman, 2011; Petersen & Berg, 2016; Berg, 2017).

### Walking speed modulation of firing rate and recruitment

To investigate the neuromuscular control of locomotor speed, we quantified speed-dependent changes in both motor unit recruitment and firing rate. We found that the majority of units were recruited more often and with larger firing rates at faster speeds (Figure 5, Figure 5–figure supplement 1). This pattern may reflect speed-dependent differences in the common input received by populations of motor neurons with varying spiking thresholds (Henneman et al., 1965). In the case of mouse locomotion, faster speeds might reflect a larger common input, increasing the recruitment probability as more neurons, particularly those that are larger and generate more force, exceed threshold for action potentials (Farina et al., 2014). This would explain our finding that motor units are more likely to be recruited during strides in which additional motor units are also recruited (Figure 7). Although this seemed consistent for each pair of motor units we measured, this was not necessarily the case given that past studies have identified motor unit substitution rather than co-activation (Westgaard & De Luca, 1999; Manning et al., 2010), potentially serving to reduce fatigue across the motor pool.

Importantly, our work only examines a subset of the movement speeds and gait patterns that mice produce. It therefore remains to be determined how rate and recruitment are reshaped as mice increase their speed up to 100 cm/s and alter coordination patterns across their limbs (trotting, bounding, etc.) (Herbin et al., 2006, 2007; Bellardita & Kiehn, 2015; Gonçalves et al., 2022). Since a majority of observed motor units, particularly in the lateral head, were already reliably recruited at the fastest speed quartile (roughly 30-40 cm/s), further speed increases might rely on either more firing rate modulation from these active units or from recruitment of more of the motor pool. Moreover, adjustments to kinematic and kinetic strategy across speeds could result from more global changes in motor unit coordination. For example, studies in drosophila (Azevedo et al., 2020) and zebrafish (Kishore et al., 2014) have demonstrated preferential recruitment of faster motor unit subtypes during rapid movements. Future studies in mice can therefore examine faster gaits to compare how different species achieve their most rapid forms of locomotion.

Considering the force production of motor units is essential to connect our observations of firing patterns to behavioral outputs. In anesthetized mice, intracellular current injections into individual motor neurons revealed that fast motor units from the triceps surae (gastrocnemius and soleus muscles of the hindlimb) reached near tetanic force at firing rates between 60-80 Hz while slow motor units reached near tetanic forces between 30-40 Hz (Manuel & Heckman, 2011). Furthermore, motor units rapidly reached these rates once active. Despite being recorded from different muscles than the ones we examined, these earlier results are relevant to our findings given that the long and lateral heads of the triceps brachii are (similarly to the gastrocnemius) biased towards fast-twitch muscle fibers (Augusto et al., 2004; Burkholder et al., 1994). Since the motor units recorded in our study had firing rates at or above the aforementioned rates immediately upon recruitment within a stride (Figure 4B), it could be that each of the units identified in this study generated near-maximal force whenever active. If units are recruited with near maximal force even at slow walking speeds, generating the additional forces needed for fast walking likely comes from recruitment of additional units. Future studies might answer this question by quantifying the force-production properties of triceps motor units during behavior, including the rapid changes in the muscle length and velocity that take place during locomotion (Edman, 1979; Gittings et al., 2012; Ting & Chiel, 2017).

Although strong correlations were observed between motor unit recruitment and limb kinematics during locomotion (Figure 6, Figure 6–figure supplement 1), it remains unclear whether such correlations actually reflect the causal contributions that those units make to limb movement. To resolve this ambiguity, future studies could use electrical or optical perturbations of muscle contraction levels (Kim et al., 2024; Lu et al., 2024; Srivastava et al., 2015, 2017) to test directly how motor unit firing patterns shape locomotor movements. The short-latency effects of patterned motor unit stimulation (Srivastava et al., 2017) could then reveal the sensitivity of behavior to changes in muscle spiking and the extent to which the same behaviors can be performed with many different motor commands.

## Methods

### Surgical implantation

All procedures described below were approved by the Emory University Institutional Animal Care and Use Committee at Emory University (IACUC protocol #201700359). Mice were anesthetized with isoflurane to implant the Myomatrix arrays. Incisions were made in the skin above the skull and above the target muscle. Forceps were used to pull the Myomatrix array through these holes so that the body of the array was entirely subcutaneous, with the Omnetics connector sitting on the skull and the array threads near the muscles. The surface of the skull was lightly scored with a scalpel and dental cement (Metabond Quick Adhesive Cement) was applied generously to fix the Omnetics connector in place and seal the opening. Myomatrix threads were then sutured (8-0 non-absorbable suture from AROSurgical) into the target muscles. Using the four threads of the customizable Myomatrix array (RF-4x8-BHS-5), we implanted a combination of muscles in each mouse, sometimes placing multiple threads within the same muscle. Threads were implanted in the triceps brachii long head and/or the triceps brachii lateral head (Table 1) and confirmed through visual inspection. We did not implant in the third (medial) head of the triceps given that it would have required an additional incision, posing more risk of surgical complications. Some mice also had threads simultaneously implanted in their ipsilateral or contralateral biceps brachii, although due to limited sample size we do not present biceps EMG data in this report. Lastly, 6-0 suture was used to close the incision. Surgeries typically took under three hours and animals were mobile shortly after removal from isoflurane.

### Behavioral methods and data collection

The treadmill used in this task had a transparent belt and base as described previously (Darmohray et al., 2019; Machado et al., 2015). A 45° angled mirror below the base allowed monitoring of side and bottom views from a single camera (FLIR Grasshopper High Performance USB 3.0 Monochrome Camera) at 330 frames per second. Separate motors controlled the left and right belts, but both were run at the same speed for every experiment. The treadmill, placed within a behavior box, was dark, with the only source of light coming from infrared light.

Experiments were conducted the day following the implant surgery up to five days post-surgery. Data presented in this study came from the first day of recording, in which signal quality tended to be highest (Chung et al., 2023). In each experimental session, mice were first briefly placed under anesthesia using isoflurane to attach a lightweight (1g) digitizing headstage (Intan RHD #C3313 16-Channel Bipolar-Input Recording Headstage) to the Omnetics connector on their skulls. Each recording session lasted approximately 45 minutes in total, and we waited at least 10 minutes after removal from isoflurane to ensure all animals were fully awake before recording began.

For five of six mice, we attempted to record 31 trials - each trial consisted of a single minute continuous running on the treadmill. The first three trials were at 10 cm/s, while the following trials were arranged in seven blocks of four trials each. Each block contained a trial at 12.5 cm/s, 17.5 cm/s, 22.5 cm/s, and 27.5 cm/s in a pseudo-random order presented identically across mice. This pseudo-random order of speeds as opposed to a strict ramping order ensured that we collected data across the full range while reducing potential effects of fatigue. For mice that became uncooperative before completing all trials, we ended the experiment early. Of these five mice, three mice completed all 31 trials, one completed 30, and the last completed 23 trials. For the sixth mouse, we again began with three trials at 10 cm/s, but only increased the speed in 2.5 cm/s increments for either two or three trials each up to a total of 14 trials. Mice were trained on their given running paradigm and habituated to the treadmill setup twice on the day before surgery. Each trial was initiated using custom Bonsai software (Lopes et al., 2015) and Arduino components to synchronize neural recordings with the camera and motor output.

We used DeepLabCut (Mathis et al., 2018) to track body parts of the mouse during locomotion (Figure 1A). We excluded points tracked with less than 90% confidence from DLC and interpolated those points from adjacent high-confidence points. The right elbow angle was estimated using markers from the shoulder, elbow, and wrist. We defined strides using the trough-to-trough minimums of the elbow angle, which occurred approximately 50-100 ms before the footstrike of the right paw. Each stride was also required to contain footstrike and liftoff of the right forepaw during forward movement. From here, we excluded strides with stride durations, stance durations, swing durations, body velocities, or body accelerations outside their respective 95% confidence interval. As a result, we kept about 80% of strides for each animal. Five of six mice had between 2600-3600 total strides, and the remaining mouse, which was run at the slower speed range, had just over 1000 strides included in analyses.

### Electromyography (EMG)

Bipolar signals from adjacent contact pairs on the Myomatrix array were extracted at 30 kHz (Intan RHD #C3100 Recording Controller) and bandpassed between 300-7500 Hz. Using up to 16 channels of high-resolution EMG from the Myomatrix arrays, motor units were identified using Kilosort 2.5, an open-source multi-channel spike sorting algorithm (Pachitariu et al., 2016). We slightly adjusted the algorithm to better fit the assumptions behind motor unit activity, including the removal of spatial decay across channels. A full description of adjustments has been previously reported (Chung et al., 2023). A total of 33 units were identified across all animals. Each unit’s isolation was verified by confirming that no more than 2% of inter-spike intervals violated a 1 ms refractory limit. Additionally, we manually reviewed cross-correlograms to ensure that each waveform was only reported as a single motor unit.

We further validated spike sorting by quantifying the stability of each unit’s waveform across time (Figure 1–figure supplement 1). First, we calculated the median waveform of each unit across every trial to capture long-term stability of motor unit waveforms. Additionally, we calculated the median waveform through the stride binned in 50 ms increments using spiking from a single trial. This second metric captures the stability of our spike sorting during the rapid changes in joint angles that occur during the burst of an individual motor unit. In doing so, we calculated each motor unit’s waveforms from the single channel in which that unit’s amplitude was largest and did not attempt to remove overlapping spikes from other units before measuring the median waveform from the data. We then calculated the correlation between a unit’s waveform over either trials or bins in which at least 30 spikes were present. The high correlation of a unit waveform over time, despite potential changes in the electrodes’ position relative to muscle geometry over the dynamic task, provides additional confidence in both the stability of our EMG recordings and the accuracy of our spike sorting.

### Data analysis

Continuous firing rates were calculated by convolving raw spike times of a motor unit with a Gaussian kernel with σ = 10 ms. This continuous result was phase-normalized across strides before calculating the mean continuous rate to identify relevant patterns such as the unit’s active duration or peak firing rate. Active duration was measured between the first and last time within a stride that the smoothed curve reached the half-height from a single spike. Overall, this method allowed for quantification of firing rates even when only a single spike was present for a stride.

### Joint model of rate and recruitment

We modeled the recruitment probability and firing rate based on empirical data to best characterize firing statistics within the stride. Particularly, this allowed for multiple solutions to explain why a motor unit would not spike within a stride. From the empirical data alone, strides with zero spikes would have been assumed to have no recruitment of a unit. However, to create a model of motor unit activity that includes both recruitment and rate, it must be possible that a recruited unit can have a firing rate of zero. To quantify the firing statistics that best represent all spiking and non-spiking patterns, we modeled recruitment probability and peak firing rate along the following piecewise function:

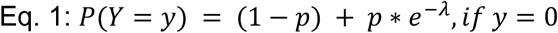

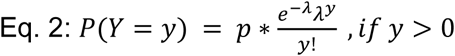

where *y* denotes the observed peak firing rate on a given stride (determined by convolving motor unit spike times with a Gaussian kernel as described above), *p* denotes the probability of recruitment, and *λ* denotes the expected peak firing rate from a Poisson distribution of outcomes. Thus, an inactive unit on a given stride may be the result of either non-recruitment or recruitment with a stochastically zero firing rate. The above equations were fit by minimizing the negative log-likelihood of the parameters given the data.

### Permutation test for joint model of rate and recruitment and type 2 regression slopes

To quantify differences in firing patterns across walking speeds, we subdivided each mouse’s total set of strides into speed quartiles and calculated rate (𝜆, Eq. 1 and 2, Figure 5A-C) and recruitment probability terms (*p*, Eq. 1 and 2, Figure 5D-F) for each unit in each speed quartile. Here we calculated the difference in both the rate and recruitment terms across the fastest and slowest speed quartiles (*p*_fast_-*p*_slow_ and 𝜆_fast_-𝜆_slow_). To test whether these model parameters were significantly different depending on locomotor speed, we developed a null model combining strides from both the fastest and slowest speed quartiles. After pooling strides from both quartiles, we randomly distributed the pooled set of strides into two groups with sample sizes equal to the original slow and fast quartiles. We then calculated the null model parameters for each new group and found the difference between like terms. To estimate the distribution of possible differences, we bootstrapped this result using 1000 random redistributions of the pooled set of strides. Following the permutation test, the 95% confidence interval of this final distribution reflects the null hypothesis of no difference between groups. Thus, the null hypothesis can be rejected if the true difference in rate or recruitment terms exceeds this confidence interval.

We followed a similar procedure to quantify cross-muscle differences in the relationship between firing parameters. For each muscle, we estimated the slope across firing parameters for each motor unit using type 2 regression. In this case, the true difference was the difference in slopes between muscles. To test the null hypothesis that there was no difference in slopes, the null model reflected the pooled set of units from both muscles. Again, slopes were calculated for 1000 random resamplings of this pooled data to estimate the 95% confidence interval.

## Acknowledgements

This work was made possible by support from NIH grants U24NS126936, R01NS109237 (SJS), NSF grant DGE-1937971 (KT), the Simons Foundation as part of the Simons-Emory International Consortium on Motor Control (SJS and MRC), the McKnight, Kavli, and Azrieli Foundations (SJS), the Portuguese Fundação para a Ciência e a Tecnologia (SFRH/BPD/119404/2016 to HGM and PTDC/MED_NEU/30890/2017 to MRC), and the European Research Council Consolidator Grant #866237 to MRC. Additionally, we thank Dr. Amanda Jacob for logistical support and William McCallum for valuable editorial comments.

---

**Figure 1-figure supplement 1.**
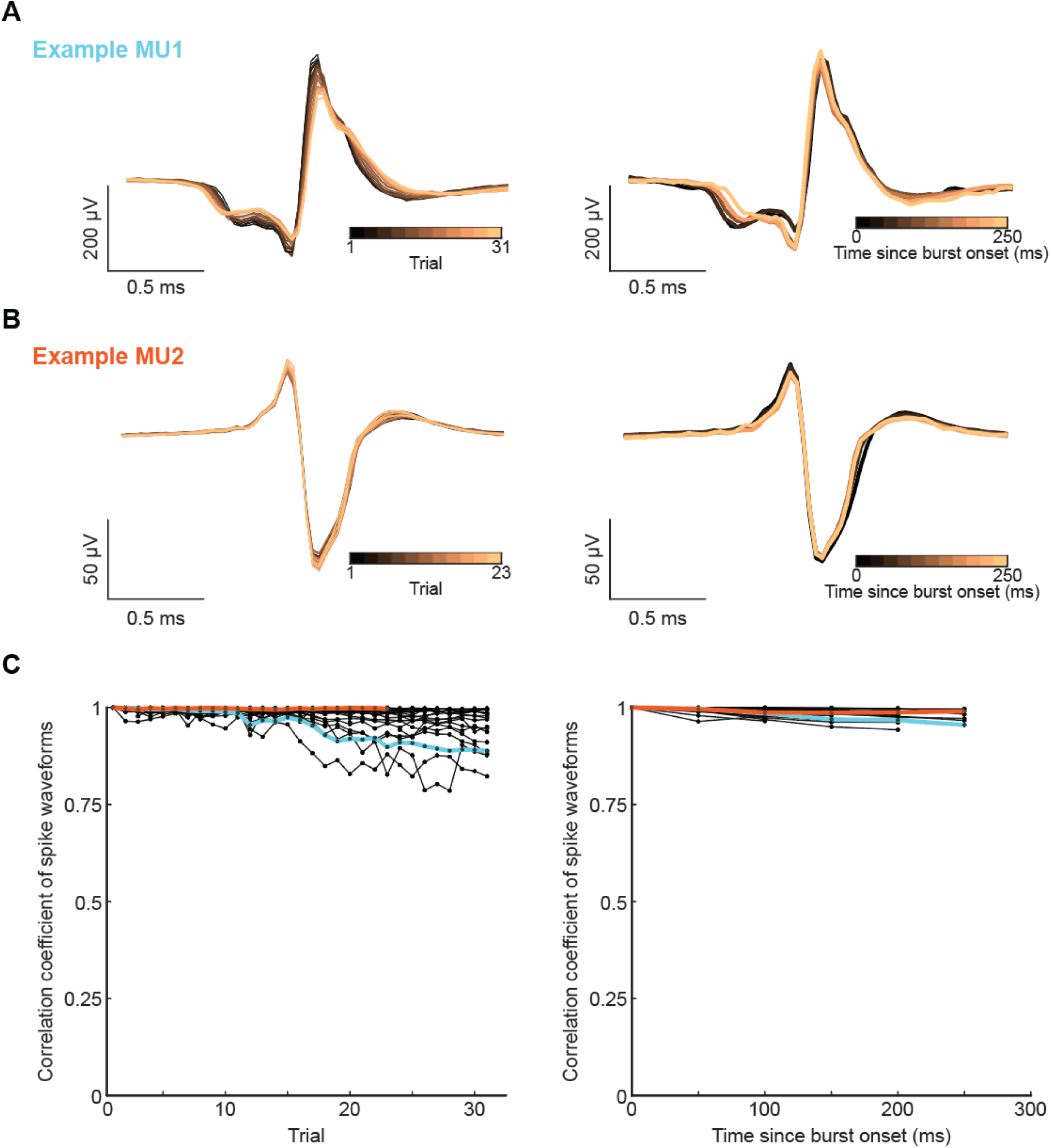
Isolated motor units had consistent waveforms. **(A,B)** Example motor unit waveforms. (Left) Median waveform calculated from a random subset of strides within each trial. (Right) Median waveform calculated from spikes binned in 50ms increments of the stride. **(C)** (Left) Auto-correlation of each unit’s median waveform between the first trial and subsequent trials. (Right) Auto-correlation of each unit’s median waveform between the first 50ms of its activity and each subsequent 50ms within the stride.

**Figure 2-figure supplement 1.**
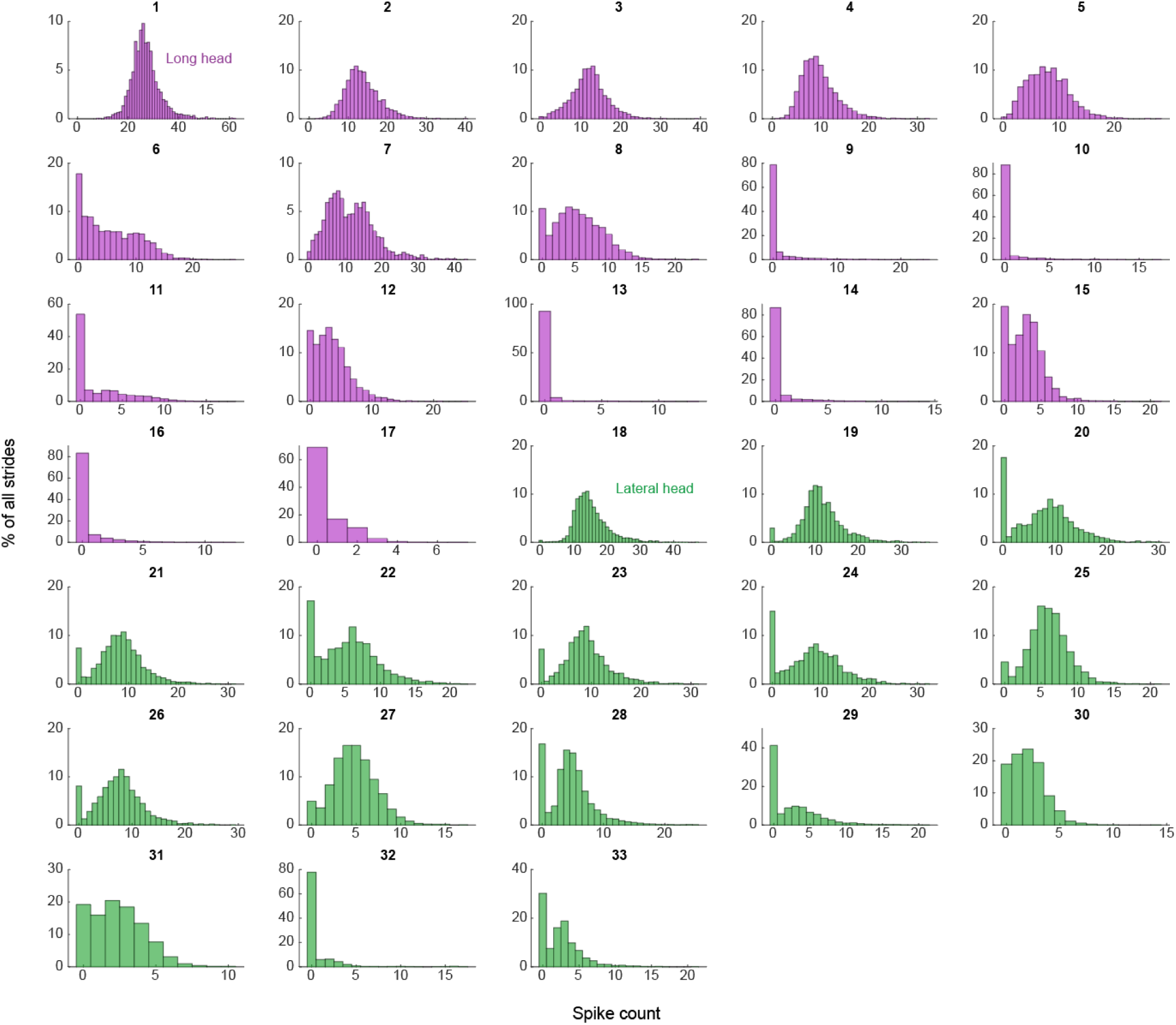
Empirical observations of spike count distributions for all units. Units are arranged sequentially to match the descending order presented in main text Figure 3. Units 1-17 are in the long head, while units 18-33 are in the lateral head.

**Figure 4-figure supplement 1.**
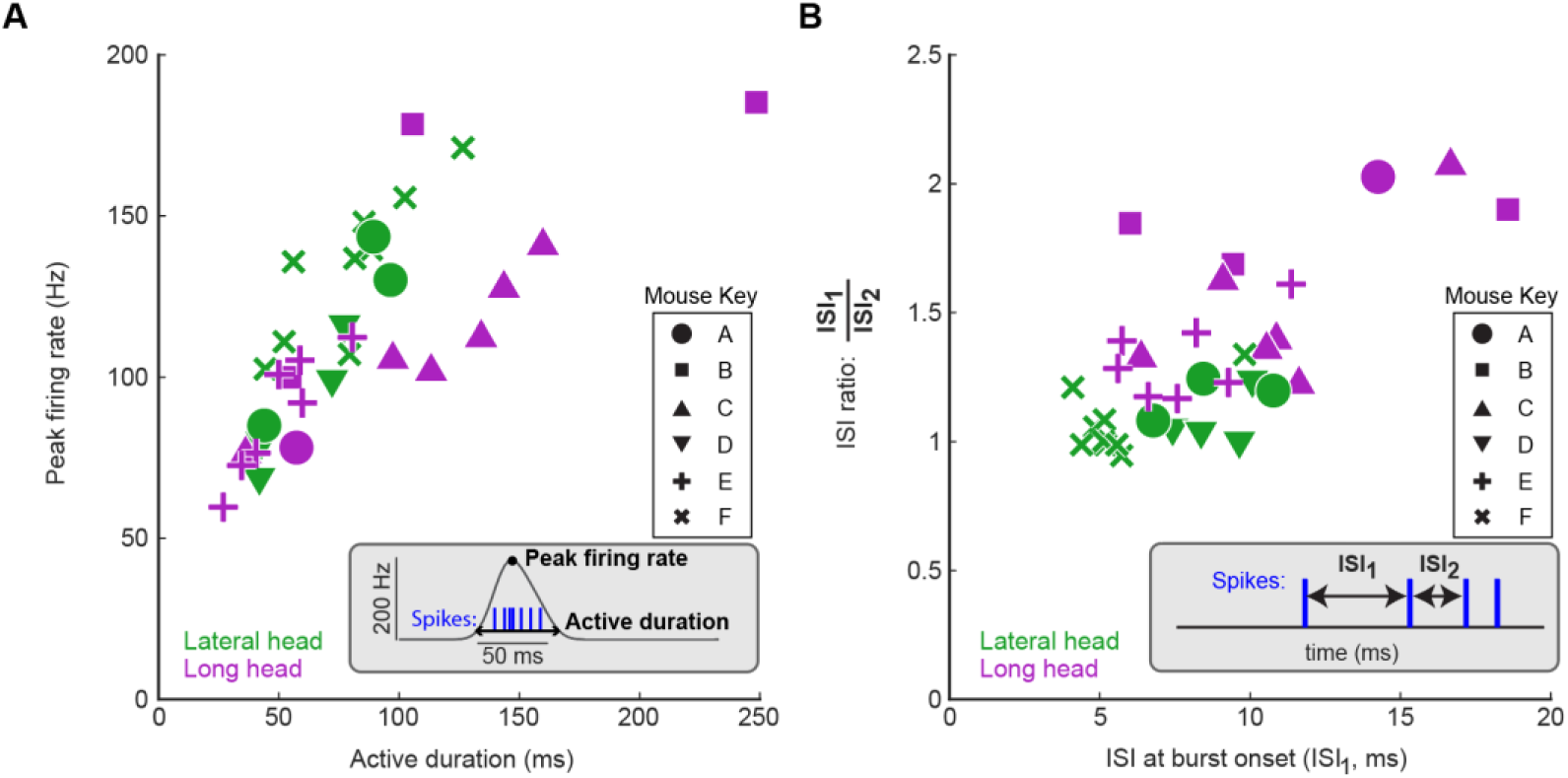
Symbols denote different animals. All other plotting conventions are the same as in Figure 4 in the main text.

**Figure 5-figure supplement 1.**
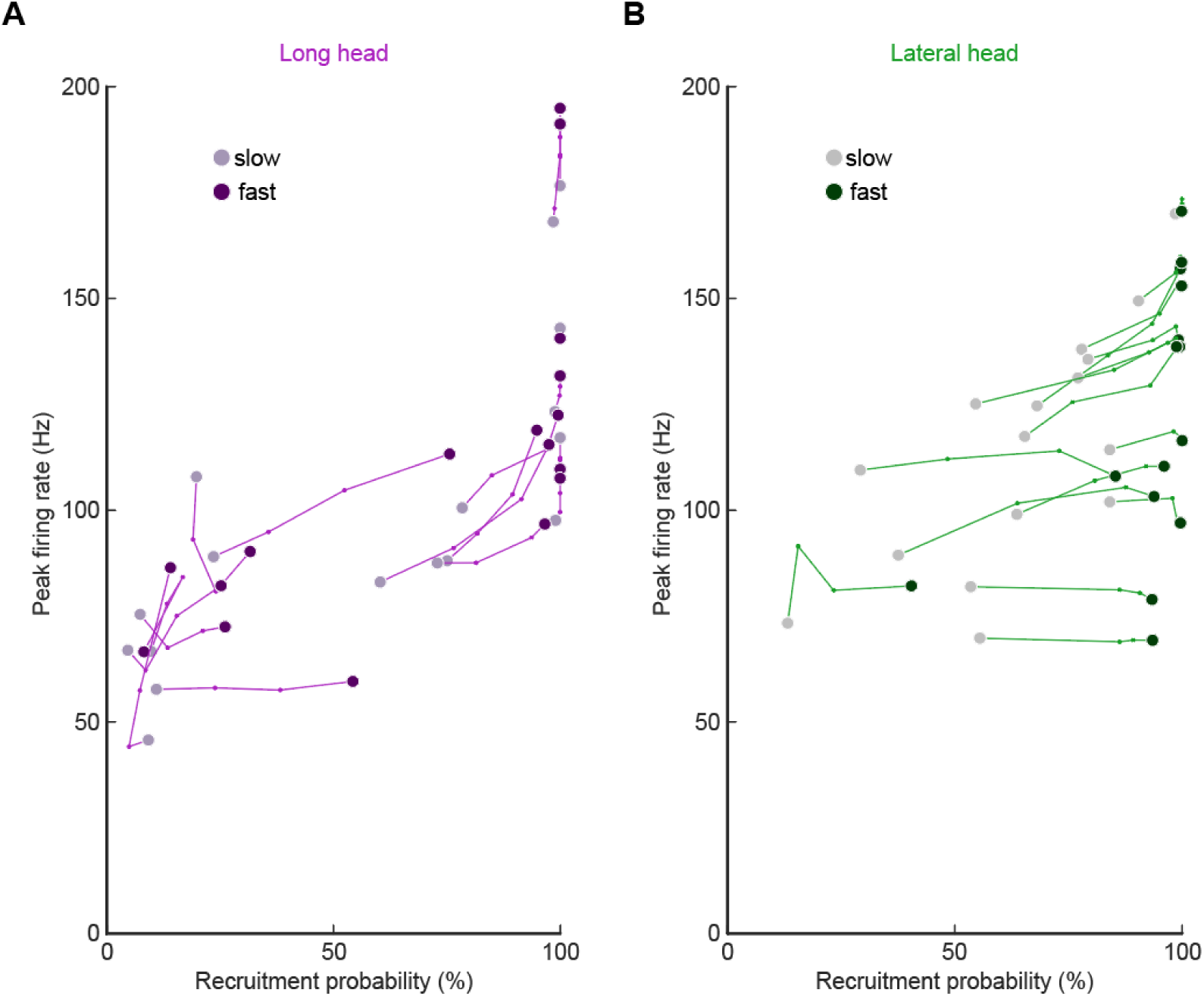
Altered firing rate and recruitment across walking speed quartiles for all motor units in the long head **(A)** and lateral head **(B)**. Each point reflects the median for the model estimate of each unit across the speeds.

**Figure 5-figure supplement 2.**
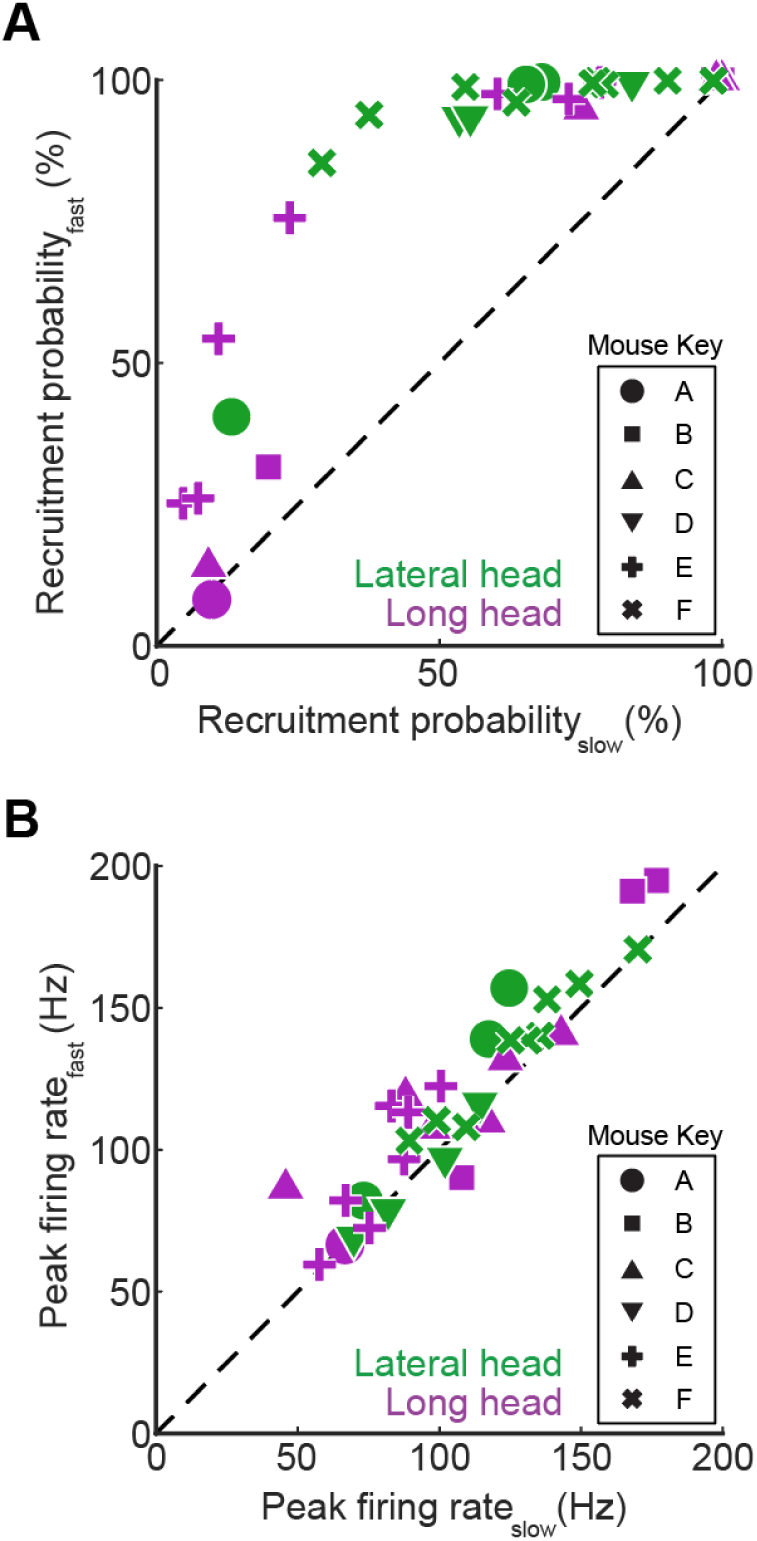
Symbols denote different animals. All other plotting conventions are the same as in Figure 5 in the main text.

**Figure 6-figure supplement 1.**
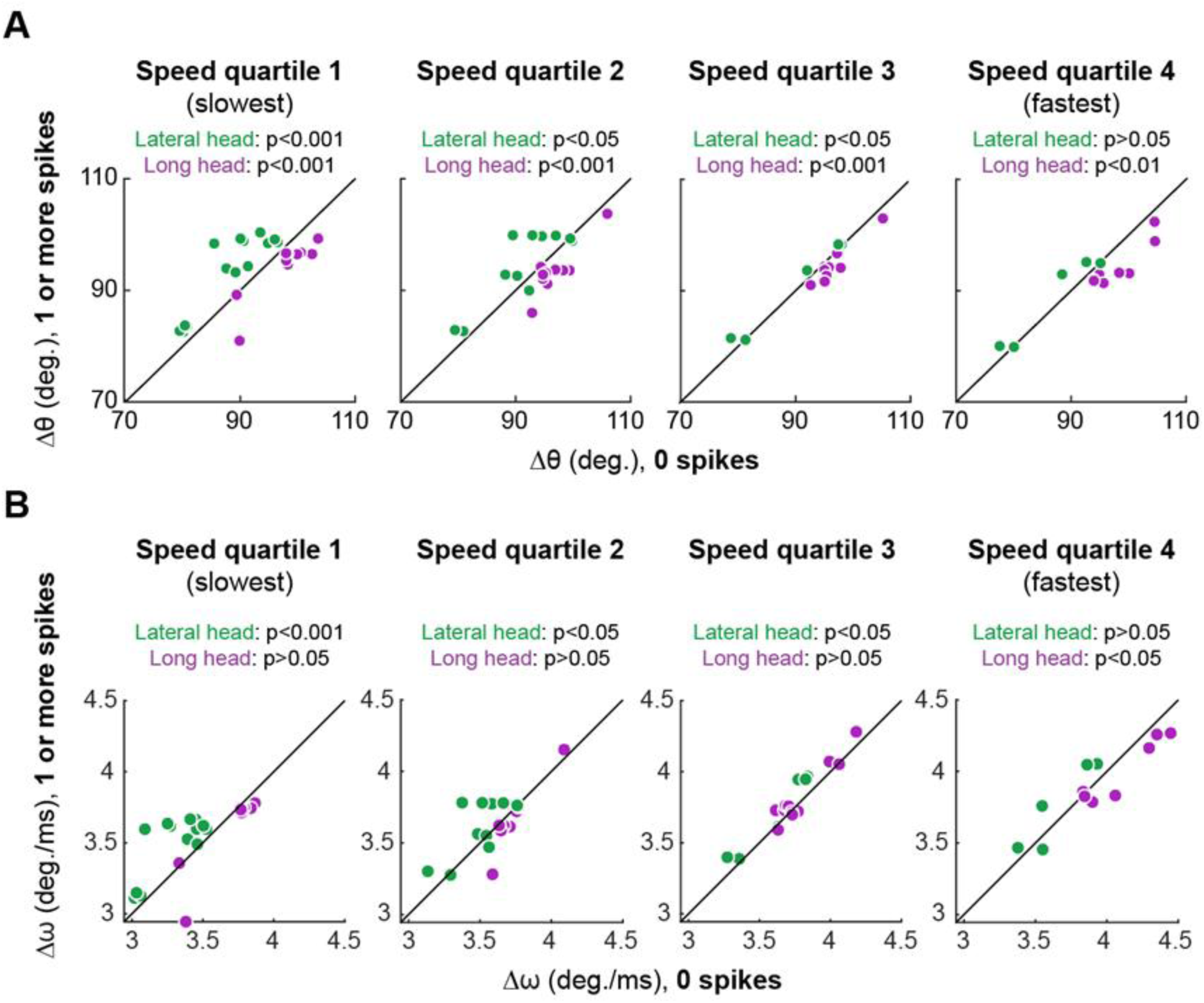
Plots show the same analysis as main text Figure 6 for each individual speed quartile of **(A)** elbow angle (Δθ) and **(B)** elbow velocity (Δω). Speed quartiles 1 and 4 are the slowest and fastest quartiles, respectively, and p-values refer to the results of Wilcoxon signed-rank tests performed separately on data from motor units of the lateral (green) and long (purple) heads of the triceps muscle. Note that most of the muscle-specific differences shown in Figure 6 (C, E) were also present when each of the four quartiles were examined individually for each muscle.

**Figure 6-figure supplement 2.**
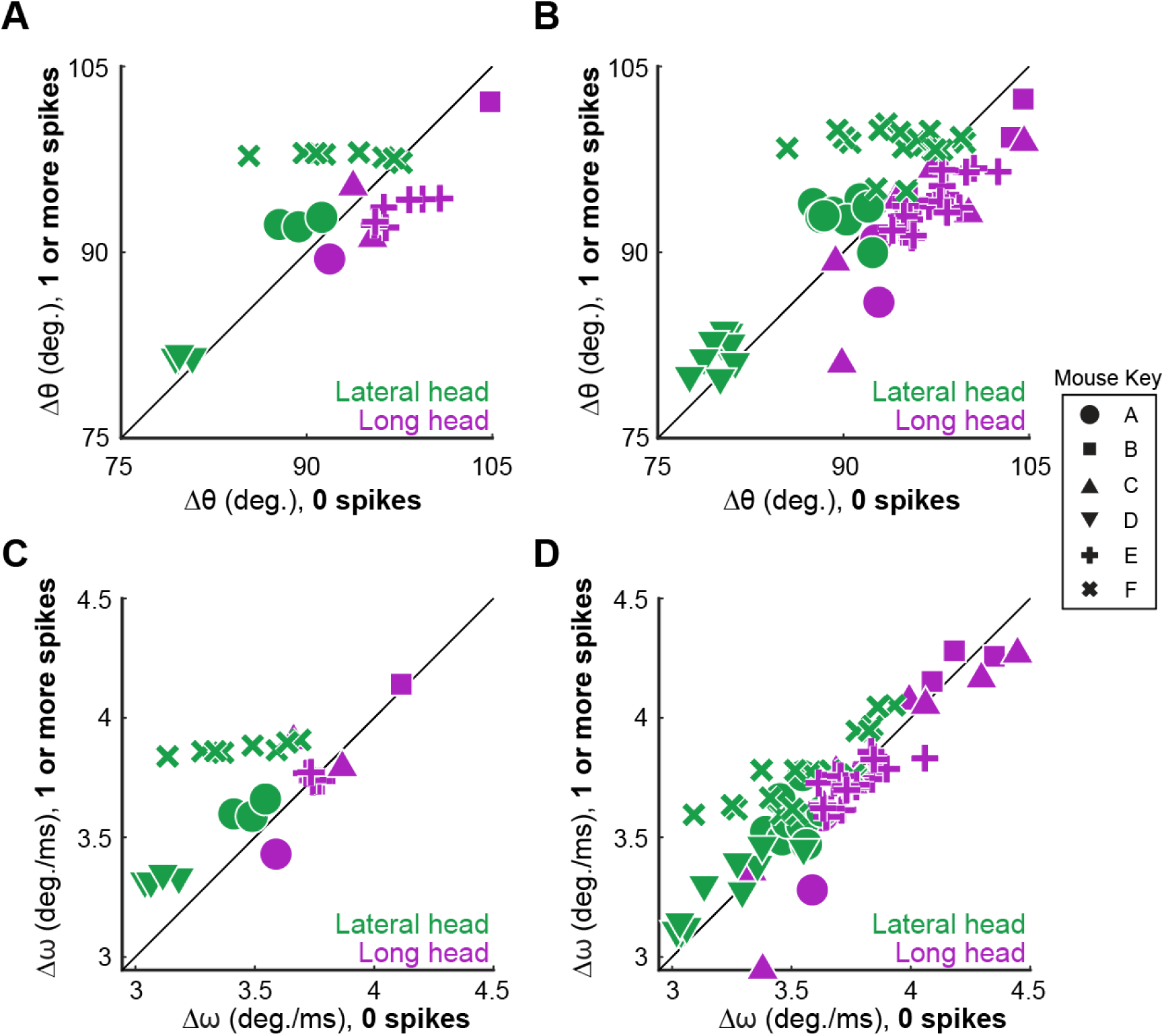
Symbols denote different animals. All other plotting conventions are the same as in Figure 6 in the main text.

